# Fluctuations in Neural Complexity During Wakefulness Relate To Conscious Level and Cognition

**DOI:** 10.1101/2021.09.23.461002

**Authors:** Pedro A.M. Mediano, Aleksi Ikkala, Rogier A. Kievit, Sridhar R. Jagannathan, Thomas F. Varley, Emmanuel A. Stamatakis, Tristan A. Bekinschtein, Daniel Bor

## Abstract

There has been considerable recent progress in measuring conscious level using neural complexity measures. For instance, such measures can reliably distinguish healthy awake from asleep subjects and vegetative state patients. However, this line of research has never explored the dynamics of conscious level during normal wakefulness. Being able to capture meaningful differences in conscious level during wakefulness may provide a vital new insight into the nature of consciousness, by demonstrating what biological, behavioural and cognitive factors relate to such differences. Here we take advantage of a large MEG and fMRI dataset of healthy adults, to examine within-subject conscious level fluctuations during resting state and tasks, by using a range of complexity measures. We first establish the validity of this approach in both neuroimaging domains by relating neural complexity measures to pre-existing techniques for capturing transitions of consciousness from full wakefulness into drowsiness and the earliest stages of sleep, finding decreased complexity as participants become increasingly drowsy. We further demonstrate that neural complexity measures in both MEG and fMRI change both within and between tasks, and relate to performance on an executive task, with higher complexity associated with better performance and faster reaction times. This approach provides a powerful new route to further explore the cognitive and neural underpinnings of consciousness.

## Introduction

Gaining a better understanding of consciousness would have profound and widespread implications for the nature of human experience, for many neurological conditions where awareness is impacted, and for a host of ethical issues. One of the greatest successes in consciousness science has been the quantification of conscious level by means of neural complexity measures.^1–4^ For instance, in a landmark paper, Casali and colleagues used transcranial magnetic stimulation (TMS) to perturb cortical activity, and then quantified the complexity in the brain’s response with electroencephalogram (EEG) via the standard measure of Lempel-Ziv complexity (LZ).^1^ This method, dubbed perturbational complexity index (PCI), was then used to distinguish between various disorders of consciousness in patients, and between healthy controls when awake versus non-rapid-eye-movement (NREM) sleep, or under general anaesthesia – with conscious states consistently associated with higher neural complexity than unconscious ones. Since this result, others have shown that the complexity of spontaneous neural activity, as measured by LZ, can distinguish between sleep states,^2^ and increases above the level of normal resting state (RS) during an altered state of consciousness due to psychoactive drugs, such as LSD.^4–6^

These results have provided a striking impetus for the continued development of consciousness science, since they are robust and highly reproducible. However, despite these successes, progress in the science of conscious level has been limited in recent years for a number of reasons. First, all the comparisons listed above concern pronounced differences between conscious states, which limits their impact in providing an explanation for the nature and possible components of consciousness during normal wakefulness. Furthermore, there is no satisfactory all-encompassing theory that explains, from first principles, the relationship between brain dynamics and conscious level. One of the main contenders, Integrated Information Theory (IIT)^7^ makes ambitious claims about the nature of consciousness, but has been criticised on conceptual^8^ and formal^9^ grounds.

An alternative approach, common in the consciousness science literature, is to search for the neural correlates of consciousness (NCC),^10^ for instance by presenting awake participants with consciously and unconsciously perceived stimuli, and investigating which regions increase in activity when stimuli are consciously detected.^11^ Although this too has yielded interesting clues as to the biological substrate of consciousness (such as the common association between increases in prefrontal parietal network activity during switches in conscious percept^12,13^), this approach also has general limitations that mean moving from the NCC to genuine mechanistic explanations for consciousness is challenging.

A new approach to the science of consciousness is therefore warranted. Here we suggest one powerful novel route that takes advantage of the recent trend towards big data to explore subtle fluctuations in conscious level during normal wakefulness, modulated by alertness, task and performance.^14–17^ In this way we are effectively combining the above two consciousness science paradigms, bridging between level- and content-centred approaches to consciousness, in a way that may yield important novel insights into the components of consciousness.^14^

We here focus on the Cambridge Centre for Ageing and Neuroscience (CamCAN) database,^18^ a large multimodal dataset which includes RS and task-based functional magnetic resonance imaging (fMRI), magnetoencelography (MEG) and behavioural data on a large cohort of healthy subjects across the adult lifespan.

First we demonstrate that neural complexity measures meaningfully reflect fluctuations in wakeful conscious level, via independent methods for assessing alertness in MEG and fMRI. Next we establish that there are clear modulations of neural complexity measures by task and performance, thus suggesting connections between consciousness and other cognitive processes. This proof of principle is the first stage in an entirely new approach to study consciousness.

## Results

### LZ fluctuates with alertness

First, we set out to investigate the relation between complexity and levels of alertness during wakefulness. As our main complexity measure we focus on Lempel-Ziv complexity (LZ), a widespread and simple yet effective complexity measure.^1–4^ In MEG, we estimate LZ as the average complexity of each individual gradiometer. For fMRI, to compensate for the low temporal resolution, we concatenate the BOLD time series from a group of regions (either the whole brain or each of Yeo’s seven networks^19^) into a single sequence, resulting in a “concatenated LZ” (LZc). To quantify alertness, we use an adapted version of the Automated Micromeasures of Alertness (AMA) algorithm^20^ for MEG, and functional-connectivity-based k-means clustering for fMRI.^21^. Following the procedures described in the Methods, we computed all complexity and alertness measures using the resting-state segments of both MEG and fMRI sessions (Fig. 1).

**Figure 1.**
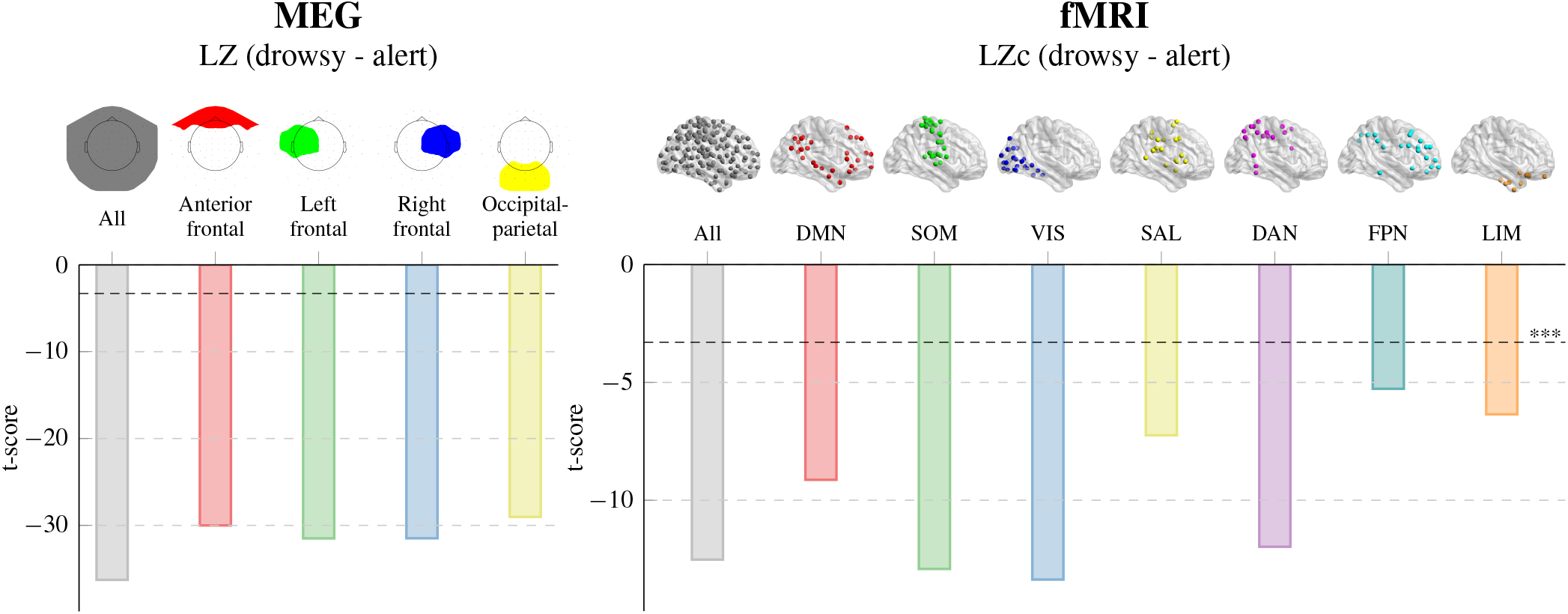
Resting-state neural complexity is reduced with drowsiness. (*left*) Effect of drowsiness on MEG signal complexity, quantified as *t*-values of linear-mixed effects models predicting LZ at different MEG sensor groups. (*right*) Two-sample *t*-values comparing drowsy subjects with alert subjects in the fMRI session. Black horizontal dashed lines show the *p* = 0.001 significance threshold. See Supplementary Figure S2 for the effect sizes of these comparisons. DMN: default mode network, SOM: somatomotor, VIS: visual, SAL: salience, DAN: dorsal attention network, FPN: frontoparietal network, LIM: limbic.

In the MEG data, we quantified the effect of drowsiness with a linear mixed-effects (LME) model predicting LZ using AMA as predictor and subject identity as random effect. The results showed a very reliable effect of drowsiness on LZ, with greater drowsiness associated with less complex brain activity (*β* = −0.0128±0.0003 bit, *t* = −35.6). It should be noted that AMA is a novel measure, not used in MEG before. Therefore we validated our results with the more traditional alpha-theta ratio measure,^22^ with qualitatively similar results (*β* = −0.0091±0.0003 bit, *t* = −26.6). Furthermore, to demonstrate that these results are not dependent on any single measure of neural complexity, we also carried out these comparisons with other measures (see Supplementary Figure S1), with again broadly similar results.

Next, we performed a similar analysis on the fMRI data, with two main differences: First, alertness level was quantified via the functional connectivity clustering method of Haimovici *et al.*^21^ instead of AMA. Second, given the much smaller number of time points in the fMRI data (5 windows of 50 TRs each), instead of using an LME we split the subjects into alert and drowsy groups, labelling as alert those who had all 5 time windows classed as awake by the Haimovici algorithm (*N* = 352), and as drowsy those who had at least 3/5 time windows classed as drowsy or early sleep (*N* = 74). As a sanity check, we verified that the proportion of awake subjects in each window decreased with time, replicating previously reported results of subjects becoming more drowsy during the scan^23^ (see Supplementary Materials).

To relate alertness and complexity, we averaged LZc across all 5 temporal windows for each subject and performed a two-sample t-test comparing the two groups. This analysis showed strong agreement with the MEG results, with drowsy subjects having significantly lower complexity than alert subjects (*t* = −12.5, *p* < 0.001). We repeated this analysis for each of Yeo’s seven brain networks^19^ in turn, and all networks showed a significant LZc reduction in drowsiness (all *p* < 0.001) to varying extents. In particular, the visual and somatomotor networks showed the lowest levels of resting LZc but the largest reductions with drowsiness, while the frontoparietal and salience network showed the smallest changes.

### LZ is modulated by task

Having established that LZ meaningfully fluctuates with alertness levels between full alertness and drowsiness, we next explored how LZ changes by task. The CamCAN database offers an excellent opportunity to explore this, thanks to its high sample numbers and wide repertoire of tasks available (see Methods). To this end, we computed whole-brain average complexity for all subjects and all tasks in both MEG and fMRI sessions (averaged across channels in MEG, concatenated in fMRI), revealing striking changes both within and between tasks (Fig. 2).

**Figure 2.**
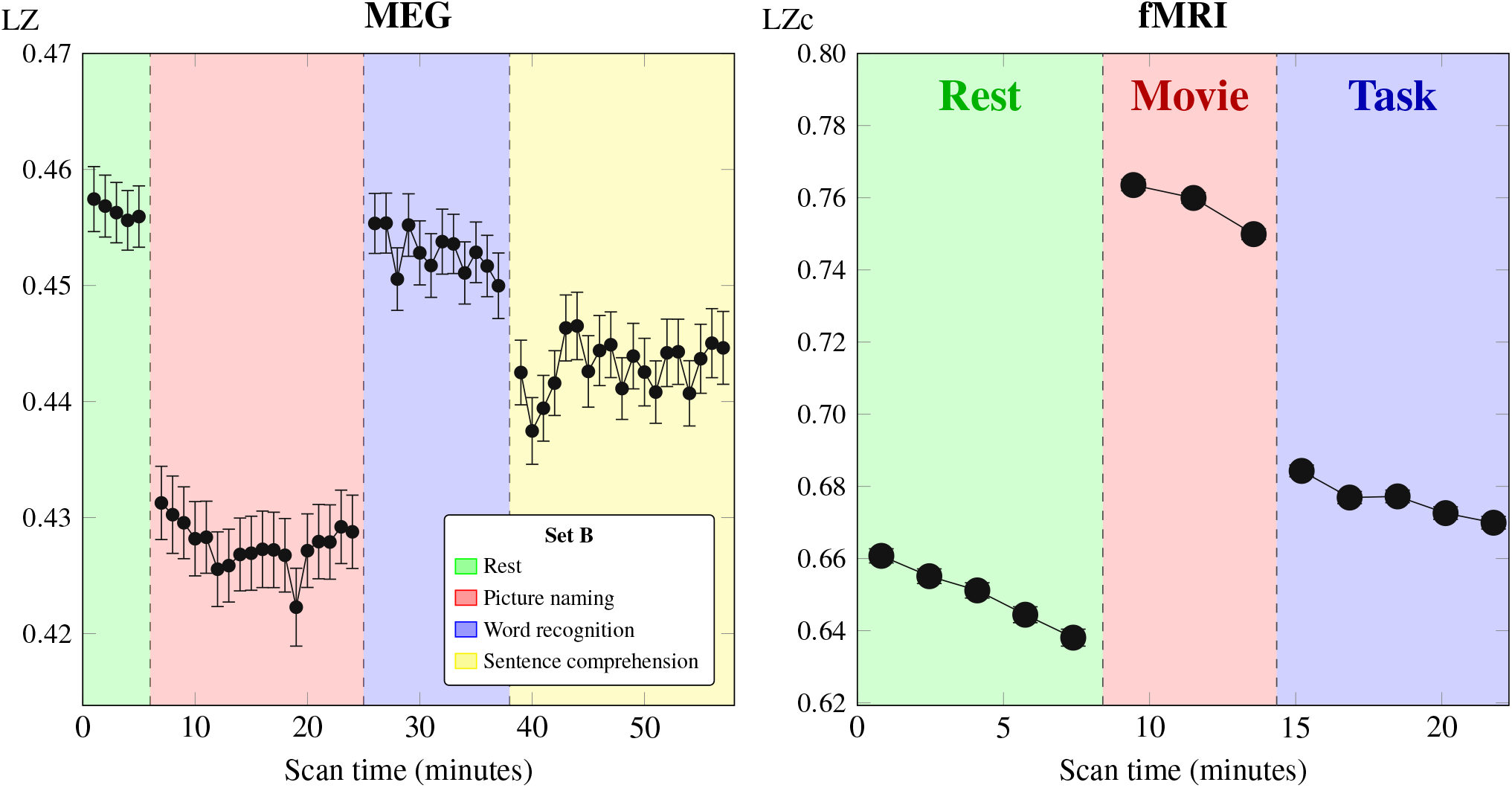
Neural complexity is modulated by task in both MEG and fMRI sessions. (*left*) Time course of average LZ throughout the MEG session, for participants in task set B (see Supp. Fig. S3 for task set A). Error bars show standard error of the mean across subjects. (*right*) Whole-brain LZc in each window of the fMRI session, showing both variation across tasks and negative drifts within tasks, suggesting subjects became progressively drowsy during each task. Note that MEG and fMRI represent different neural processes and LZc is different from LZ, so values are not directly comparable between the two.

In the MEG session, participants showed a wide spectrum of LZ values, which varied markedly between tasks (see Supplementary Figure S5). Of all 12 pairwise comparisons between tasks (including sets A and B), 11 of them had significant LZ differences at the *p* < 0.001 level using one-sample t-tests, with the only non-significant comparison being between multi-mismatch negativity and incidental memory in task set A. Interestingly, all active tasks (i.e. tasks that required subjects’ input; including SNG, picture naming, and SC) had lower complexity than wakeful rest, suggesting that more cognitively demanding tasks in fact *decrease* neural complexity. This result shows that LZ, and other common measures of conscious level (see Supplementary Material), at least with the high temporal resolution of M/EEG, are also sensitive to the neural dynamics of cognitive processes, opening a new fertile scientific ground at the intersection of these two phenomena.^14^

As in the previous section, the fMRI results were largely in agreement with the MEG results. Between tasks, there was a significant difference between the three conditions for the whole brain (all *p* < 0.001), and within each task LZc declined steadily for all tasks (including resting state), possibly reflecting a general drop in alertness or interest in the task. In line with previous results,^4^ LZc increased with respect to the resting-state baseline when participants passively watched a movie, providing further evidence that contextual factors can affect subjects’ spontaneous neural complexity. Note that, however, in fMRI LZc is higher during task than in wakeful rest, unlike the results for the MEG data. The cause of this discrepancy is unclear, but it is unlikely to be caused by the concatenation of channels in LZc since it also occurs with average channel-wise LZ (see Supplementary Figure S4).

Finally, it is worth mentioning a few important differences in LZc between brain networks as they relate to the tasks in the fMRI session. For both task and movie-watching, the visual and somatosensory networks showed the largest increases with respect to rest, while the frontoparietal network showed slightly weaker task distinctions compared to the other networks. Additionally, the dorsal attention network was the only one to show a positive interaction effect between task and movie-watching, with higher complexity during task (*t* = 6.77, *p* < 0.001; see Supplementary Figure S6).

### LZ correlates with task performance

The fact that these complexity measures track alertness and are modulated by task suggests they can be used to explore the connection between consciousness and cognition. For this we focused on the one attentionally demanding, executive task present in both fMRI and MEG task sets: the SNG task.

In each trial of the SNG task, subjects are presented with one of four types of visual stimuli. For two of them (go-left and go-right signals) the subject must respond via button press as quickly as possible (with their left or right hand, respectively). For the other two stimuli (stop and no-go signals), the subject must withhold their response and wait for the next trial. For this analysis we focused on the reaction times in the go trials and the total number of omission and commission errors, regardless of which stimulus type they were triggered by (see Supplementary Figure S7).

In the MEG session, we analysed the trial-by-trial relationship between LZ and reaction time using an LME model to predict reaction time in a given trial, using the LZ of the preceding 4 s as predictor and subject identity as random effect. This revealed a small but consistent negative effect (*β* = −533± 50 ms bit^−1^, *t* = −9.13), showing that subjects responded faster in trials with a higher level of pre-stimulus LZ. To investigate this further, we analysed whether subjects with higher inter-trial LZ variability also had higher RT variability. In agreement with the previous result, we found a significant positive correlation between the coefficient of variation (CV) of reaction time and the CV of LZ (*r* = 0.31, *p* = 0.023), indicating that subjects whose LZ fluctuated more widely during the session also saw their reaction times fluctuate. In the fMRI session, although trial-by-trial data is not available (due to the limited temporal resolution of fMRI), meaning we couldn’t perform the same analysis as in MEG, we also found evidence of the relationship between LZ and task performance: average LZc was anti-correlated with the subject’s total number of errors, including omission and commission (*r* = −0.36, *p* = 0.017).

Taken together, these results suggest that in the cognitive domain neural complexity is characteristic of a state of conscious readiness, in which neural dynamics switch between a diverse set of possibilities, able to adapt quickly in response to task demands. More generally, they reinforce our main thesis that LZ can be used to mechanistically connect consciousness and cognition: higher cognitive demands may call on the same (or similar) neural processes that underpin consciousness, which can be measured via LZ.

## Discussion

### Fine-grained fluctuations in consciousness and complexity

In this paper we have demonstrated, using both MEG and fMRI, that neural complexity measures can be used to track fluctuating conscious level during wakefulness. Furthermore, these same complexity measures change for different tasks, reflecting changes in the subject’s condition and cognitive state. Finally, we have shown that these complexity measures correlate with accuracy and reaction time in an executive task. These results suggest that conscious levels not only change from awake to sleep states, but fluctuate, in line with alertness and drowsiness, on a moment by moment basis – and these spontaneous fluctuations offer us an opportunity to investigate the inner workings of consciousness during wakefulness.

In addition to the temporal dynamical aspects captured by LZ, this type of analysis allows us to explore the spatial heterogeneity of complexity throughout the brain. Although there were no differences between local regions in the MEG data, in fMRI there were intriguing differences between the seven Yeo brain networks. For example, the somatomotor and visual networks had lower complexity overall, both during alertness and drowsiness. In contrast, the frontoparietal network had one of the highest LZc overall, demonstrated the least decline from alert to drowsy states, and was less modulated by task than other networks. One explanation for this is that the frontoparietal network, by having the highest LZc generally, even sustained during drowsiness, is more centrally involved in supporting conscious contents than lower-level networks involved in sensory processing or motor output (which would be closely in accord with the existing literature^11,13^).

### Connecting neural complexity and cognitive processing

On the cognitive level, our results demonstrate that complexity measures are predictive of behaviour and cognitive performance. We demonstrated this in fMRI using broad temporal averages, as well as in MEG with a trial-by-trial analysis, showing in both cases that higher neural complexity was associated with better performance (in terms of fewer errors and faster reaction times). Although the posited association between executive processing and consciousness is not new,^24^ here we’ve provided, to our knowledge, the first neurally-driven evidence supporting its connection, albeit in a provisional form we hope that we, and others, will build on in future studies.

Interestingly, however, there is one important aspect for which the MEG and fMRI results disagree: in the MEG session LZ was lower for all tasks than during resting state, while the opposite was true in fMRI. To the best of our knowledge this is the first reported discrepancy between LZ changes across multiple imaging modalities, with previous works showing consistent results in fMRI, MEG, and EEG.^2,4,5,25^ However, it is worth noting that these reported consistent changes all concern drastic changes in conscious state, such as between wakeful rest and deep sleep or anaesthesia. Thus, the fine-grained structure of the fluctuations of consciousness during wakefulness studied in this paper poses a new challenge for empirical tests of current theories of consciousness and cognition.

Nonetheless, there are also a few consistent patterns across fMRI and both MEG sessions that are worth mentioning. Most notably, both fMRI and MEG passive tasks (i.e. multi-mismatch, word recognition, and movie watching) induce consistently higher complexity than active tasks. This could be caused by these tasks calling on different underlying cognitive mechanisms, although without further experiments we cannot discard an effect of salience driving the LZ results: for example, the word recognition task in MEG had unexpected non-words, and the movie shown in the fMRI session was engaging and fast-paced, while the active tasks were generally repetitive – which could explain the differences in complexity between the two.

Overall, although future work is needed to explain these results, we can attempt to interpret them within the framework of the Entropic Brain Hypothesis (EBH).^26^ Put briefly, the EBH states that the richness of phenomenal experience should be accompanied by a similarly rich repertoire of neural dynamics, quantifiable via complexity or entropy measures.^27^ If, as the EBH postulates, LZ essentially captures the variety of phenomenological events, and given the cognitive tasks were relatively prescribed and uniform compared to resting state, it is natural that entropy is higher in resting state. At the same time, this variety of phenomenology is more likely to be captured by the higher temporal resolution of MEG, but not fMRI, which only captures slower neural processes due to the timescale of the haemodynamic response^28^. Overall, this discrepancy highlights the need for future work exploring the profile of neural complexity across scales, from aggregated cellular activity to BOLD signals, and paints a more nuanced picture regarding the interpretation and physiological relevance of LZ deserving further investigation.

### Limitations and future work

Our approach in this study has been openly empirically-driven, applying an experimentally validated set of measures to a large neuroimaging dataset and analysing the results. Nonetheless, like previous studies linking complexity measures to consciousness,^1–3^ a theoretical foundation behind these analyses is largely underspecified. For instance, it is not clear how LZ (and its variants) map onto elements of candidate theories of consciousness, such as the differentiation or integration components of IIT^1^ – or, for that matter, whether the different measures proposed historically by IIT can in fact meaningfully assess integration and differentiation in brain dynamics,^30,31^ and how these relate to cognition.^32^ Therefore, we need better theories to fully interpret these results and incorporate them into a broader picture of consciousness as a neurobiological phenomenon. In this sense, we see these results as a more challenging testbed for upcoming theories of consciousness, that are willing to go beyond gross changes in conscious level (such as between wakefulness and sleep or coma) and able to explain these subtle fluctuations of consciousness during wakefulness.

One concern with the analyses presented here is whether there may be some specific details of LZ that generate these results. To rule out this possibility, we have used a range of measures, including multiple flavours of LZ, multi-scale entropy,^33^ and context-tree weighted predictors^34^ (see Supplementary Material). Broadly speaking, all these measures behaved similarly, suggesting that the link between neural complexity and fluctuating conscious levels during wakefulness and between tasks is a robust finding independent of any individual measure. However, complexity is a fundamentally multi-dimensional concept,^35^ and there are plenty of complexity measures that differ heavily from LZ on conceptual grounds.^2^ Furthermore, all the measures explored here quantify the temporal statistics of individual regions, ignoring possible high-order statistical structures taking place across regions.^31,39^ This study opens the door to future work investigating the subtle differences between other types of complexity measures and their specific relationships with consciousness and cognition.

Finally, a separate issue concerns whether the observed changes in neural complexity measures are driven by changes in spectral power. For instance, in MEG, it could have been the case that as participants became more drowsy, and their faster alpha power diminished while their slower theta power increased, this spectral shift automatically supported less diverse activity patterns, thereby lowering the measured LZ. Although preliminary results on the MEG SNG data suggest that some, though not all, complexity changes are not explained by spectral power,^40^ this analysis should be extended to all aspects covered here (as well as to previous studies with larger changes in conscious state). Overall, we see this line of work as a necessary step to better understand the dynamical and mechanistic basis of neural complexity measures, so we may better leverage them in empirical and theoretical studies of consciousness.

### Final remarks

For these analyses we leveraged the power of big data via the CamCAN^18,41^ database, which allowed us to resolve subtle changes in the measures of interest thanks to its large size, and make first steps towards elucidating the components of consciousness. We believe that this will be a powerful, flexible new approach to accelerate progress in consciousness science generally.

Although the results presented here provide intriguing putative clues as to the components of consciousness, we believe that their main purpose is to demonstrate the utility of this approach for future, more directed, studies than was possible with the CamCAN dataset. We envisage these future studies to focus on fluctuations in complexity measures during wakefulness and these changes to be linked to specific cognitive components. In this way, a taxonomy of the potential cognitive machinery of conscious could be found.

## Methods

### Participants

We examined a pre-existing dataset, collected by the Cambridge Centre for Ageing and Neuroscience (CamCAN) project, involving adults who had undergone both fMRI and MEG at two stages, approximately 2 years apart. In the first stage, here termed CamCAN650, 650 adult participants (after attrition for technical issues) underwent a series of fMRI and MEG RS and task-based scans. In the second stage, here termed CamCAN280, 280 of the CamCAN650 participants returned for more extensive fMRI and MEG scans. These scanned subjects were behaviourally tested to ensure they were neurologically normal. Ages ranged from 18 to 88, with a roughly equal-*N* split between deciles. This age range was designed to explore to the effects of age on cognition and the brain. Although we also found modulatory effects of age on some of the effects reported below, these are not relevant to the core results and are described elsewhere (paper in preparation). For more details on the CamCAN database, see Shafto *et al.*^18^ and the CamCAN website.^3^

### Tasks

For both MEG and fMRI, in addition to RS, the experiment included different sets of tasks of varying cognitive load. Here we analysed fMRI and MEG data for all tasks using the complexity measures described below. Furthermore, some of them were active tasks (i.e. that included behavioural responses) that were centrally cognitive, which allowed us to perform additional analyses linking complexity values to performance, under the putative assumption that consciousness is related to cognition, given its involvement in most high-level cognitive skills.^11,42^

The tasks are described in more detail elsewhere,^18,41^ and are briefly summarised here for completeness. See the Supplementary Material for further information about tasks in both MEG and fMRI sessions.

#### MEG

Given that the CamCAN650 session did not include cognitive tasks, here we focus on the CamCAN280 session. Participants were divided into two groups (of approximately *N* = 140 each). All participants first underwent a 5 min resting state scan and then carried out a set of tasks, which was different for groups A and B (details in Table S1 and the original CamCAN publications^18,41^). Here we analysed complexity values for all tasks, and additionally analysed the Stop-Signal Go/No-Go task (SNG) (the only core cognitive task presented) for connections to task performance.

#### fMRI

CamCAN650 included resting state, passive movie watching, and a basic sensorimotor task, where participants pressed a button if they heard or saw a stimulus. Here we analyse all participants in CamCAN650 for the RS and movie sessions, and focus on CamCAN280 to link complexity values to performance on executive control tasks. In particular, we analyse the sub-group of subjects within CamCAN280 that took the same SNG as in the MEG session, to enable a comparison across scanning modalities.

### Neural complexity and alertness measures

#### MEG

The main elements of our MEG (and fMRI) data analysis fall into two categories: previously validated measures of alertness (which we apply only to RS data), and measures of complexity (which we apply to all data).

The first alertness measure we use is adapted from the Automated Micromeasures of Alertness (AMA) algorithm, a machine-learning-based technique which classifies EEG data into alert, drowsy, or various sleep stages.^20^ In order to work with MEG data, we swapped the key EEG electrodes the original algorithm uses with equivalent MEG channels, and, given that the CamCAN data is unlikely to include any significant portions of actual sleep, removed the parts of the algorithm that classify sleep stages. The result is a classification of each epoch as “alert” or “drowsy” which is objective, non-relative, and can be compared between subjects.

Given the novelty of this method, we also used another measure of alertness: the ratio of alpha to theta power (Alpha-Theta Ratio; ATR), a robust and empirically well-established marker of drowsiness in eyes-closed EEG.^22,43^. We took theta power in the 3–5 Hz range, alpha power in the 8–12 Hz range, and calculated the ratio between these two frequency bands as the mean of all MEG gradiometer sensors per epoch. More drowsy epochs were those where alpha power was reduced and theta power was increased.

In terms of neural complexity measures, in line with recent literature^1–4^ we focused on Lempel-Ziv complexity (LZ)^44^ as a measure of neural signal diversity. In short, the procedure to estimate LZ from a time series of neural activity of length *T* is as follows: first, the signal is binarised around its median. Then it is scanned sequentially using the algorithm by Kaspar and Schuster,^45^ counting the number of different “patterns” in the signal. Finally, following Ziv^46^ this number is normalised by log_2_(*T*)*/T* to yield an estimate of the signal’s entropy rate.^47^ This process is repeated for all channels and the results averaged into a single whole-brain average LZ.

Although our focus here is on using LZ, other suitable complexity measures exist, some of which have already been successfully applied to distinguish between conscious levels.^3,48,49^ Therefore, to demonstrate the robustness of our findings we replicate our analyses with a range of entropy-based complexity measures, and show they yield consistent results (see Supplementary Material).

#### fMRI

As with the MEG analysis, we apply two types of measures to the fMRI data: a measure of alertness for resting-state fMRI, and several measures of complexity.

To measure fluctuations in alertness during the fMRI session, we relied on a study by Haimovici and colleagues that combined simultaneous EEG and fMRI while subjects transitioned from full alertness to deep sleep.^21^ Haimovici *et al.* used a k-means algorithm to cluster subjects’ functional connectivity into “awake” and “sleep” clusters. For our analysis, we took the cluster centroids from the Haimovici study in order to classify the CamCAN data into high- and low-alertness segments (of 100 s each), roughly equating to the “awake” and “drowsy” micromeasures classification in the MEG dataset.^4^

To measure complexity we again focus on LZ complexity, for consistency with the MEG analysis. It is worth noting that although fMRI has less temporal resolution than MEG, recent studies^25^ have shown that fMRI LZ is as robust of an index of conscious level as it is in other modalities. Nevertheless, given the short length of fMRI BOLD time series (compared to MEG), instead of computing LZ on each region separately, we concatenated all time series into a single one-dimensional signal, and then computed and normalised LZ as described in the MEG section, resulting in a “concatenated LZ” (LZc). This procedure was performed for all parcels in the Schaefer 300-region atlas, as well as in those subsets of parcels that belong to each of Yeo’s seven networks.^19^ Again, as in the MEG analysis we verified the robustness of our results through other similar measures of complexity (see Supplementary Material).

### Data preprocessing

#### MEG

All data were automatically analysed with Matlab scripts using the SPM12^50^ and EEGLAB^51^ libraries. First, Independent Components Analysis (ICA) was used to reduce the effects of eye blinks. The data was split into gradiometer and magnetometer channels, and only the 204 gradiometer channels were selected for further preprocessing, due to superior signal-to-noise ratios. Following this, the data was downsampled to 250 Hz, and filtered with a 0.5–30 Hz bandpass filter. Data were epoched to 4 s segments in both RS and task recordings. For a further analysis relating task performance with complexity at the trial level, we focused on the only CamCAN executive task (SNG), and extracted 1 s epochs immediately prior to each trial. Epochs with muscle artefacts (widespread frequency above 120 Hz across channels) or unusually reduced signal (less than 25% of average signal compared with the rest of the session) were excluded from the analysis.

#### fMRI

Functional MRI data were preprocessed in two separate pipelines. The first one uses the AAL atlas^52^ and replicates the preprocessing steps of Haimovici *et al.*^21^ in order to use their functional-connectivity-based drowsiness classification. The second pipeline uses the Schaefer 300-region parcellation grouped into Yeo’s seven networks,^19^ and was used for all complexity analyses. Note that the usage of two separate pipelines is necessary for a network-based complexity analysis, since Yeo’s networks are defined on the cortical surface, and mixing surface- and volume-based parcellations (like the Schaefer and AAL parcellations, respectively) reduces the accuracy of spatial localisation.^53^ The details of both pipelines are described in the Supplementary Material.

## Acknowledgements

We thank Ariel Haimovici and Enzo Tagliazucchi for assistance with the drowsiness classification algorithm, and Darren Price for comments on earlier versions of this work. P.M., A.I. and D.B. are funded by the Wellcome Trust (grant no. 210920/Z/18/Z). R.K. was supported by the UK Medical Research Council SUAG/047 G101400 and a RadboudUMC Hypatia Fellowship.

## Author contributions statement

D.B. conceived the experiments, preprocessed and analysed the results, and co-wrote the manuscript; P.M. analysed the results and co-wrote the manuscript; A.I. preprocessed and analysed the results and co-wrote the manuscript; R.K. conceptually contributed to the manuscript; S.R.J. conceptually contributed to the manuscript, assisted in the preprocessing and analysed the results; and T.A.B. contributed conceptually to and co-wrote the manuscript. All authors reviewed the manuscript.

## Footnotes

1 Although there is some preliminary evidence in toy logic-gate models,^29^ it is unclear how or to what extent these results apply to whole-brain dynamics.

2 Examples include measures of information synergy^36^ fractal dimension,^25^ metastability^37^ or network connectivity,^38^ to name a few.

3 http://www.mrc-cbu.cam.ac.uk/datasets/camcan/

4 Note that the Haimovici study labelled the non-alert state as “sleep” – however, given that in the CamCAN650 dataset subjects still responded regularly on the sensorimotor task when classed in this state, we believe that “drowsy” is a more appropriate label than sleep. See Supplementary Material for details.

