## Supplementary Material for "Fluctuations in Neural Complexity During Wakefulness Relate To Conscious Level and Cognition"

*{pam83, db246}@cam.ac.uk

#### fMRI preprocessing pipelines

##### First pipeline

Full details of this preprocessing pipeline are found in the original CamCAN publications.[1,2] Preprocessing was performed with SPM12 software using the Automatic Analysis[3] batching system. Data were unwarped using field-map images, realigned to correct for motion, slice-time corrected, co-registered to the T1 image, normalised into MNI space, and finally smoothed with a 12 mm Gaussian kernel.

We extracted BOLD time series using the AAL parcellation.[4] Then, as in Haimovici *et al.*,[5] six head motion realignment parameters (translation and rotation), and their three first derivatives, were regressed out, and time series were bandpass filtered in the 0.01–0.1 Hz range. However, unlike Haimovici *et al.*, we did not regress physiological noise signals out.

Subjects with excessive movement during scanning were excluded based on two criteria: too much wavelet de-spiking (for the first [CamCAN650] stage data only), or too high root mean square of volume-to-volume displacement. As the amount of de-spiking is quantified by a percentage of voxels containing a spike in a given volume, subjects with an average spike percentage of two standard deviations above the group average were excluded. Volume-to-volume displacement was estimated with the approach of Jenkinson *et al.*,[6] and here also a value of two standard deviations above the group average led to exclusion. Furthermore, a subject was excluded if there were any flat or near-flat signals (standard deviation of a BOLD signal < 0.5 and mean value < 50 before motion regression) as this suggested a poor overlap between the AAL atlas and functional images.

##### Second pipeline

In the second pipeline data were preprocessed with FreeSurfer's standard preprocessing tools, which included motion correction, slice-timing correction, and intensity normalisation.

We extracted BOLD time series in FreeSurfer using Schaefer's 2018 300-ROI parcellation.[7] Then, as suggested by Power *et al.*[8] we regressed head motion realignment parameters with a Volterra expansion (translation and rotation parameters, current and immediately preceding timepoint, squares of both timepoints[9]). Finally, as in the first pipeline, time series were bandpass filtered in the 0.01–0.1 Hz range.

Subjects with poor functional-anatomical registration (`tkregister-sess` QA value > 0.6) were excluded from further analysis. Also, similarly to the first pipeline, we excluded any subjects that had flat or near flat signals (standard deviation of a BOLD signal < 0.1 after motion regression), or if RMS framewise displacement[10] was more than 1.5*IQR above the 75$^{\text{th}}$ percentile of group values, where IQR stands for the 25–75 inter-quartile range.

#### Additional complexity measures

As mentioned in the main text, in this article we have focused on LZ as our main complexity measure, emphasising its interpretation as a measure of unpredictability (more technically, entropy rate) of a stochastic process. This is of course not the only notion of complexity that exists,[11] although we argue it is a particularly useful one in the context of physiological time series analysis. In this spirit, we validated our main analyses with the following complexity measures, all based around the concept of predictability and its operationalisation through Shannon entropy.

With these ideas in mind, we chose the following additional complexity measures:

- Entropy rate computed with a Context Tree Weighted (CTW) predictor,[12] a Bayesian variable-order Markov model able to capture long-term temporal structure in discrete sequences.
- Multi-Scale Entropy (MSE),[13] a measure of complexity at multiple temporal scales based on sample entropy.

- For completeness we also use (un-concatenated) LZ in the fMRI data; although, as mentioned in the main text, BOLD time series are too short for an accurate estimation of LZ, so in this context we expect this measure to be less sensitive than LZc.

#### Evidence the fMRI "sleep" state is actually a drowsiness state

Although the resting state and movie watching CamCAN650 fMRI conditions were passive, the sensorimotor task was active, where participants responded with a finger press to any stimulus. This task, albeit simple, allowed us to explore whether the 100 s windows labelled as "sleep" should be considered as sleep or drowsiness: If the sleep state genuinely reflected times when participants were asleep, then there should be a marked absence of response in those time windows. If instead accuracy and reaction times were only marginally affected during these non-alert time windows, then this would be evidence that the "sleep" stages should more appropriately be classed as "drowsy." Most participants of those who had clean fMRI data on this task ($N = 535$) were "awake" in all five 100 s time. Only nine participants had the majority of time windows (3 out of 5 or more) classed as sleep/drowsy during the task. Although such a small sample is not appropriate to carry out statistical tests on, Table S1 clearly shows that the sleep/drowsy time window should be classed as "drowsy." Across the 120 trials, even the sleep/drowsy group is still close to ceiling, with just 1.5% lower accuracy than the fully awake group. However, the sleep/drowsy group is approximately 60 ms slower and more variable in RT, compared to the fully awake group. This pattern of slower, more variable RT is common in drowsy performance.[14,15] Therefore, these results suggest that the "sleep" label in this dataset is better conceptualised as drowsiness.

**Table S1.** Behavioural results on the CamCAN650 sensorimotor task, split by alertness levels.

| **Participant type** | **5/5 time windows "Awake"** | **$\geq$ 3/5 time windows "sleep/drowsy"** |
|---|---|---|
| **Mean % correct** | 99.5 | 98.0 |
| **Mean RT (ms)** | 308 | 371 |
| **Coefficient of variation RT** | 0.211 | 0.267 |

### Supplementary figures and tables

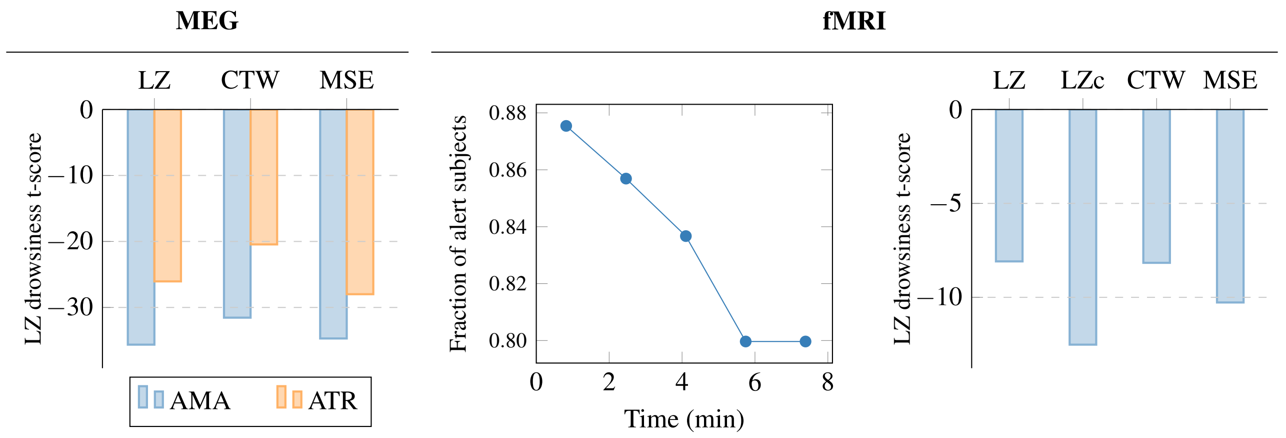


**Figure S1.** **Supplementary results on the relation between complexity and alertness**. (*left*) T-scores of drowsiness LME coefficients for various whole-brain average neural complexity measures, calculated for the resting-state (RS) MEG session. (*middle*) Fraction of alert subjects during the 5 windows (of 50 TRs each) of the RS fMRI session, as computed with Haimovici *et al.*'s[5] clustering algorithm. (*right*) Two-sample t-scores between alert and drowsy subjects for various neural complexity measures averaged throughout the whole RS fMRI session. AMA: automated micromeasures of alertness, ATR: alpha-theta ratio.

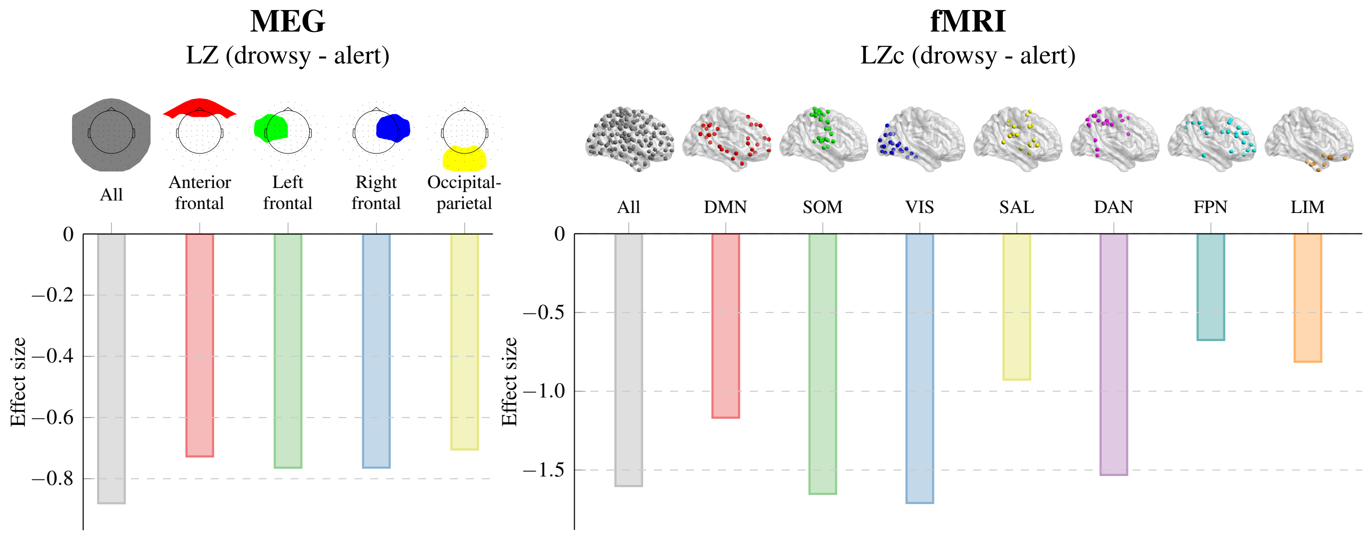


**Figure S2.** **Effect sizes for drowsiness-induced complexity reduction in resting-state data**. All comparisons and regions of interest are taken as in Figure 1 in the main text, but using effect size (Cohen's $d$) instead of $t$-values. For MEG $d$ is computed as suggested by Feingold,[16] taking the regression coefficient divided by the residuals' standard deviation. For fMRI $d$ is computed with the standard formula, as the difference in means divided by the pooled standard deviation.

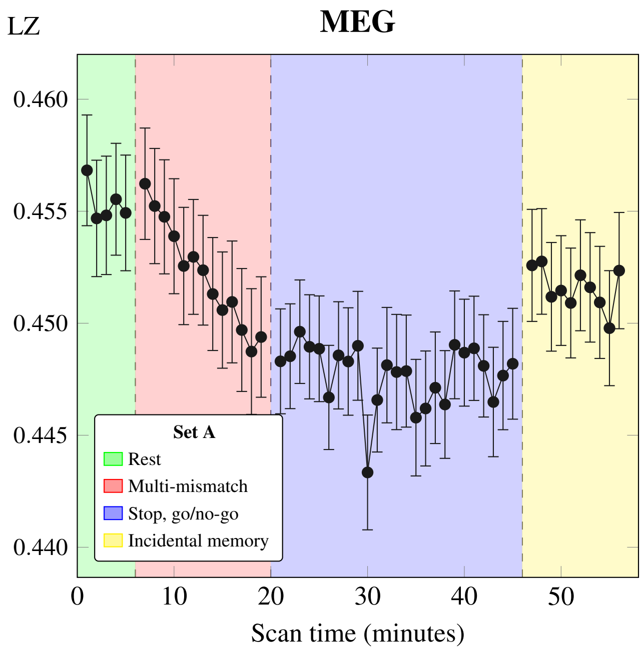


**Figure S3. Temporal profile of LZ complexity in MEG task set A**. See Table S2 for task details. As seen for task set B, complexity fluctuates noticeably between tasks. Interestingly, within-task complexity decreases with time in the multi-mismatch task, the only passive task in the set.

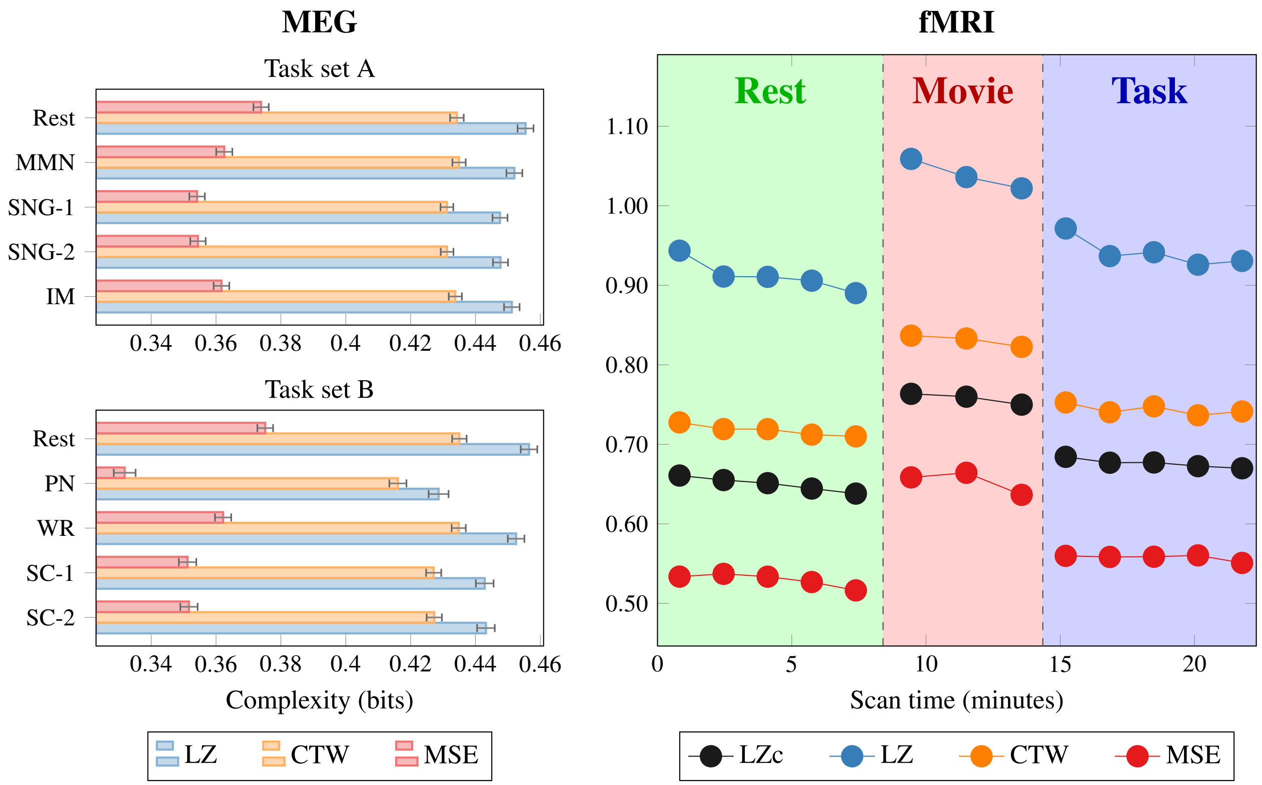


**Figure S4. Results on the relation between complexity and task replicated with alternative complexity measures**. See Tables S2 and S3 for task details. MMN: multi-mismatch negativity, SNG: stop signal go/no-go, IM: incidental memory, PN: picture naming, WR: word recognition, SC: sentence comprehension.

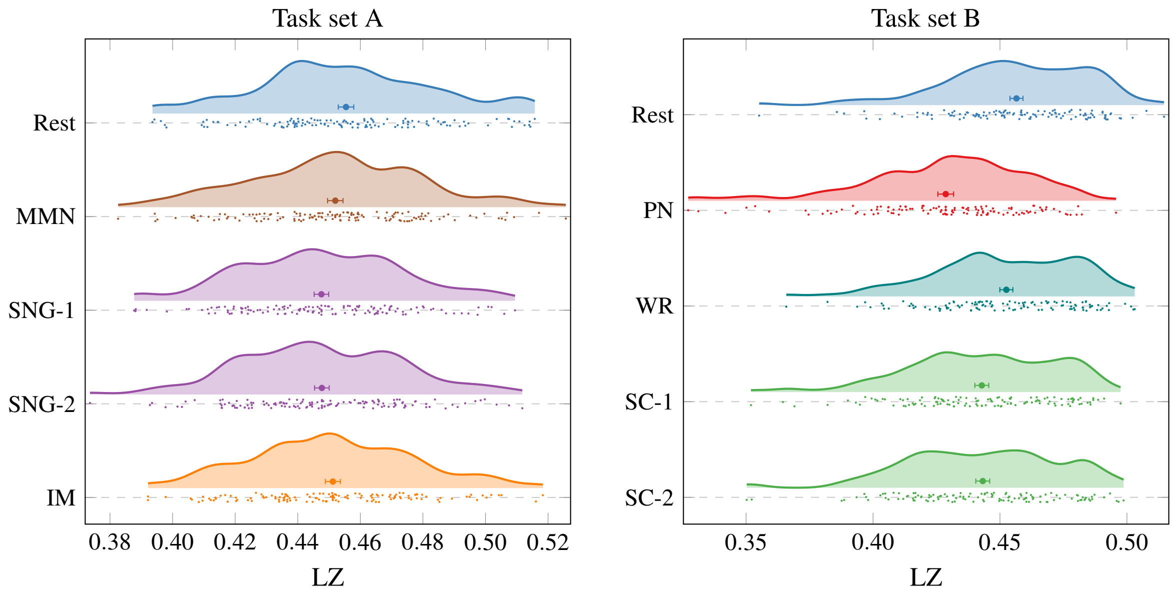


**Figure S5. Raincloud plots showing the distribution of whole-brain LZ across all subjects in both MEG task sets**. Error bar represents standard error of the mean. See Tables S2 and S3 for task details. MMN: multi-mismatch negativity, SNG: stop signal go/no-go, IM: incidental memory, PN: picture naming, WR: word recognition, SC: sentence comprehension.

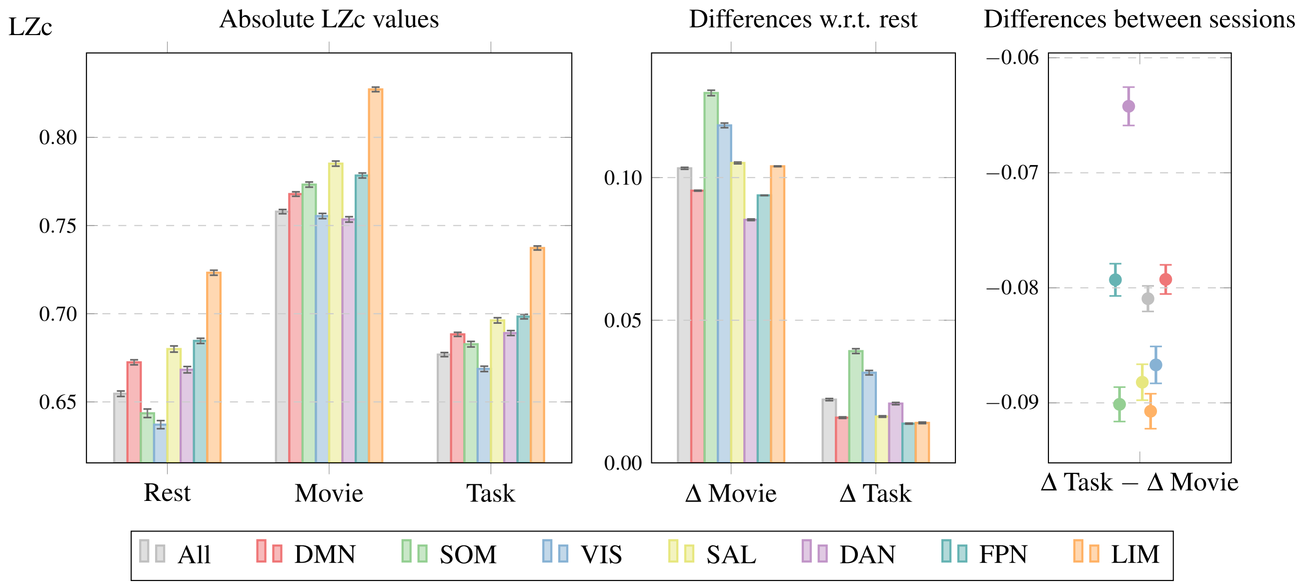


**Figure S6. Differential effects of task on neural complexity across brain networks**. (*left*) Absolute LZc values for the seven Yeo brain networks[7] for each segment of the fMRI session, showing increased complexity in all networks during movie watching and sensorimotor task. (*middle*) Difference in LZc with respect to rest. The visual (VIS) and sensorimotor (SOM) networks showed highest differences between tasks. (*right*) Difference between task-induced and movie-induced LZc changes w.r.t. rest, showing the dorsal attention network (DAN) had comparatively higher LZc in the task. Error bars represent standard error of the mean across subjects.

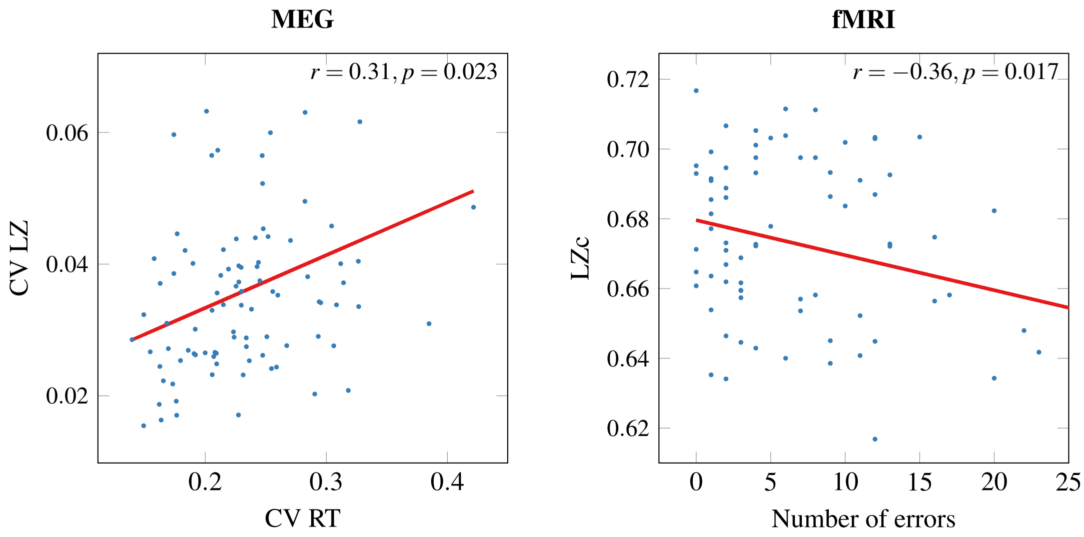


**Figure S7. Neural complexity correlates with cognitive performance in the stop-signal go/no-go (SNG) task**. (*left*) In MEG, the coefficient of variation (CV) of LZ correlates with the CV of subjects' reaction time, with higher LZ leading to lower reaction times (as determined with an LME model; see text for details). (*right*) In fMRI, a lower average LZ is associated with a higher number of errors.

**Table S2.** List of behavioural tasks that participants carried out during the CamCAN280 MEG sessions that we analysed here. The task analysed additionally for performance-complexity measure associations is underlined. Half of subjects were in group A and half in group B.

| Task name (group A or B) | Task | Main contrast / MEG analysis | Active / passive | Eyes open / closed |
|---|---|---|---|---|
| Resting State (A & B) | None | None | Passive | Closed |
| Multi-Mismatch (A) | A series of auditory tones are presented, with standard tones intermixed with various kinds of "deviant" tones. The participant merely has to listen to the tones (i.e. no active task). | Standard vs deviant tones. | Passive | Open |
| Stop-Signal, Go/No-Go (A) | If there is a black arrow pointing to left or right, participants are required to press left/right buttons. If the black arrow changes to red (stop-signal) or is initially presented in red (no-go), then participants should not make any responses. | Go vs stop-signal vs no-go, as well as correct (omission of response) vs incorrect (response made incorrectly) stop-signal and no-go trials. | Active | Open |
| Incidental Memory (A) | View complex pictures, and respond to those (rare) stimuli that contain a moon. | Initial vs repeat viewing of same picture. | Active | Open |
| Picture Naming (B) | Participants name aloud one of 302 different object pictures as fast as possible. | Correct vs incorrect naming. | Active | Open |
| Word Recognition (B) | Visually presented root and suffix of a compound word or non-word. No task (just passive reading). | Real complex words (e.g. "farmer") vs pseudo complex words (e.g. "corner") vs non-words (e.g. "goated") vs words with only a root (e.g. "scandal") vs words with neither a root nor suffix (e.g. "biscuit") vs basic non-word consonant strings. | Passive | Open |
| Sentence Comprehension (B) | Unambiguous and ambiguous sentences are read aloud, with a pause before the final word, which is either classed (by a button response) as acceptable or unacceptable. | Examination of the activation and resolution of syntactic ambiguity. | Active | Open |

**Table S3.** List of behavioural tasks that participants carried out during the CamCAN650 and CamCAN280 fMRI sessions that we analysed. Half of subjects during the 2nd stage were in group A or B. Then half of subjects were in group C or D.

| Task Name (Stage [650, 280] and Group [A-D]) | Task | Main contrast / fMRI analysis | Active / passive | Eyes open / closed |
|---|---|---|---|---|
| Resting State (CamCAN650 and CamCAN280C & CamCAN280D) | None | None | Passive | Closed |
| Movie Watching (CamCAN650) | Watch an 8 minute black and white drama TV show. | Differential responsiveness to different sections of the story. | Passive | Open |
| Sensorimotor task (CamCAN650) | Visual, auditory auditori-visual stimuli presented. Participants respond with a finger press to any stimulus. | Stimulus type and responsiveness. | Active | Open |
| Fluid Intelligence (CamCAN280A and CamCAN280B) | Four patterns are presented, in easy and hard versions, and participants have to choose the “odd one out” via a button press. | Hard vs easy blocks, correct vs incorrect trials. | Active | Open |
| Stop-Signal Go/No-Go (CamCAN280A) | If there is a black arrow pointing to left or right, participants are required to press left/right buttons. If the black arrow changes to red (stop-signal) or is initially presented in red (no-go), then participants should not make any responses. | Go vs stop-signal vs no-go, as well as correct (omission of response) vs incorrect (response made incorrectly) stop-signal and no-go trials. | Active | Open |
| Visual Short-Term Memory (CamCAN280D) | One, two, or three displays of coloured dots are shown per trial, with each display rotating. Following an 8s delay, participants have to move the probe display (whose colour matches one of the previous displays) to the direction that the previous corresponding stimuli were moving. | Short term memory capacity (performance), either within or across set size. | Active | Open |
